# Synchronization of oscillatory growth prepares fungal hyphae for fusion

**DOI:** 10.1101/2022.09.25.509415

**Authors:** Valentin Wernet, Vojtech Kumpost, Ralf Mikut, Lennart Hilbert, Reinhard Fischer

**Author notes:** corresponding author, Web: www.iab.kit.edu.

## Abstract

Communication is crucial for organismic interactions, from bacteria, to fungi, to humans. Humans may use the visual sense to monitor the environment before starting acoustic interactions. In comparison, fungi lack a visual system, instead, hyphae use a cell-to-cell dialogue based on secreted signaling molecules to orchestrate cell fusion and establish hyphal networks. Hyphae alternate roles as signal-sender and signal-receiver, as can be visualized via the putative signaling protein, Soft, which is recruited in an oscillatory manner to the respective cytoplasmic membrane of interacting hyphae. Here, we show that signal oscillations already occur in single hyphae of *Arthrobotrys flagrans* in the absence of a potential fusion partner. They occurred in the same phase as growth oscillations. Once two fusion partners came into each other’s vicinity, their oscillation frequencies slowed down (entrainment phase) and transit into anti-phasic synchronization of the two cells’ oscillations with frequencies of 130 +/-20 sec. Single-cell oscillations, transient entrainment, and anti-phasic oscillations were reproduced by a mathematical model where nearby hyphae can absorb and secrete a limited molecular signaling component into a shared extra-cellular space. We show that intracellular Ca^2+^ concentrations oscillate in two approaching hyphae, and depletion of Ca^2+^ in the surrounding affected vesicle-driven extension of the hyphal tip, abolished single-cell molecular oscillations and the anti-phasic synchronization of two hyphae. Our results suggest that single hyphae engage in a “monologue” that may be used for exploration of the environment and can dynamically shift their extra-cellular signaling systems into a “dialogue” to initiate hyphal fusion.

**Significance statement:** Communication at the cellular level often relies on chemical signal exchange. One prominent example is the fusion of fungal hyphae to form complex hyphal networks. As opposed to mating-type dependent cell fusion, cell-fusion events described here occur in genetically identical cells. Relying only on one chemical signaling channel raises the question of how communication is initiated. We discovered that individual hyphae constantly perform signal oscillations, comparable to a cellular “monologue” until they meet another hypha with which they then coordinate signal oscillations in a cell-to-cell dialogue. We also show that signal oscillations are mechanistically interlinked with calcium-dependent growth oscillations. Although the signaling molecule(s) has not been identified yet, it is highly likely linked to the hyphal growth machinery.

## Results and Discussion

Oscillations are common phenomena in biology with periods from seconds to hours, to days, to years (1-3). In fungi, oscillations are a well-described feature of hyphal tip extension, where calcium ions control vesicle accumulation, actin depolymerization, and vesicle fusion with the tip membrane (4). In addition, signal oscillations have been described at the hyphal tips during the fusion of hyphal cells (5-7). These signal oscillations were named *cell-to-cell dialogue* and are probably based on a conserved diffusible signaling molecule, which remains to be discovered (8-10). As the cell-to-cell dialogue appears to be based on a single, chemical communication channel, it is proposed that the two involved cells undergo oscillatory secretion of a signaling compound with a refractory period that prevents self-stimulation and can contribute to overcoming critical threshold concentrations (1). We recently studied the cell-to-cell dialogue during trap formation in the nematode-trapping fungus *Arthrobotrys flagrans* (*Duddingtonia flagrans*) (9). *A. flagrans* is able to switch from a saprotrophic to a predatory lifestyle, and sophisticated signaling regimes between the fungus and its prey, *Caenorhabditis elegans*, control trap formation, attraction of the prey and attack of the nematode (11-14). One of the hallmark proteins during the cell-to-cell dialogue is the *soft* protein, SofT, which shows oscillatory recruitment to the plasma membrane of interacting cells (7, 9). Although its molecular function remains elusive, SofT is thought to be involved in generating a signal during the cell-to-cell dialogue. It is essential for cell fusion in filamentous fungi and acts as a scaffold protein of the cell wall integrity pathway (15-18). An open question is if and how cell-to-cell communication is activated once a hyphal fusion partner appears in the vicinity. To address this question, we monitored SofT-GFP in *A. flagrans* growing on low-nutrient agar (LNA) and observed oscillatory recruitment of SofT to single hyphal tips with a mean period of 130 ± 20 (SD) seconds (n=50 in 5 hyphae) without other hyphae in their vicinity **(Fig. 1a, d, e, Movie S1)**. These dynamics resembled oscillating tip growth described in *Aspergillus nidulans* and raised the question whether and how these two processes might be connected in *A. flagrans* (4). Therefore, we monitored the mCherry-tagged orthologue of *A. nidulans* chitin synthase B (DFL_009443) in *A. flagrans*, which acts as a cargo marker for intracellular transport and is crucial for polarized growth (19, 20). The fluorescent fusion protein oscillated at growing hyphal tips with a mean period of 125 ± 37 (SD) seconds (n=50 in 7 hyphae) **(Fig. 1b, d, e, Movie S2)**. To compare the oscillations of SofT and ChsB, we created a strain with both proteins labelled. SofT-GFP localized at the same time at growing tips as mCherry-ChsB **(Fig. 1e, Movie S3)**. However, mCherry-ChsB showed a wider frequency distribution and shorter periods. Thus, the localization of mCherry-ChsB appears to be independent of SofT **(Fig. 1f)**. Indeed, localization and oscillation of mCherry-ChsB at the tip were not affected by deletion of *sofT* **(Fig. 1f)**. Deletion of *chsB* in *A. nidulans* is detrimental for hyphal growth and was therefore not further analyzed in *A. flagrans* (21, 22). Our results show that hyphae are constantly sending signals in a form of constant “self-talk”, or monologue, possibly to explore the environment for a fusion partner.

**Fig. 1:**
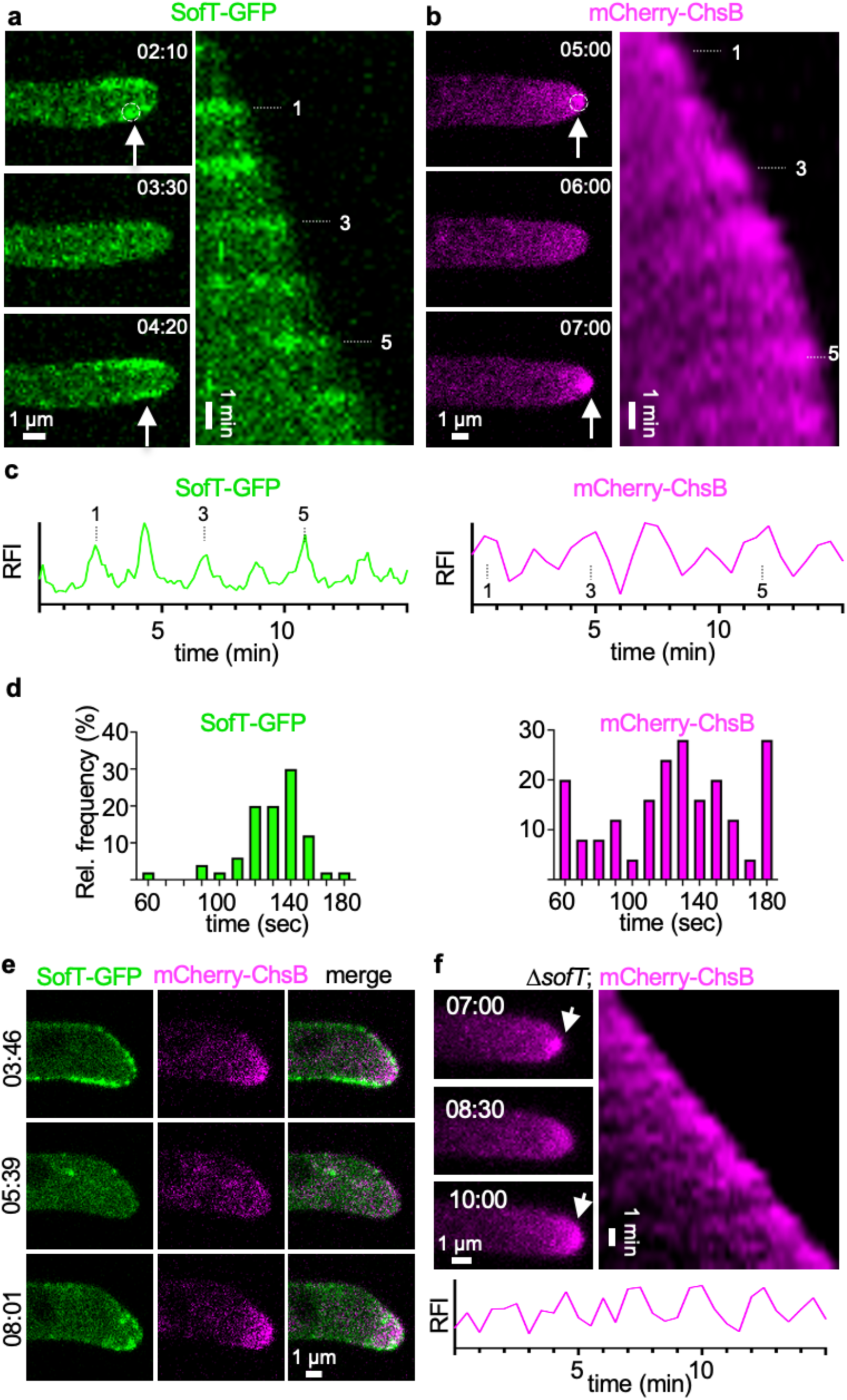
Oscillatory recruitment of signaling proteins during hyphal growth of *A. flagrans*. **(a, b)** Time course of SofT-GFP and mCherry-ChsB localization at hyphal tips. Arrows indicate the accumulation of the proteins at the hyphal tips. Dotted circles indicate an area of 6×6 pixels which was used to measure the fluorescent intensity over time depicted in (c) of GFP-SofT at the lower edge of the plasma membrane or of mCherry-ChsB at the apex. (b) is a maximum-intensity projection of the time-lapse sequence. A kymograph was created for each time course drawing a line (pixel width 5) along the growth axis of the respective hypha. Numbers indicate the count of oscillating accumulation of each fusion protein at the hyphal times during the time course. **(c)** Relative fluorescent intensity (RFI, y-axis, arbitrary units) at the hyphal tips of a – b was measured over the time course (x-axis in minutes). **(d)** The interval between two peaks at hyphal tips was counted and depicted as relative frequency (y-axis) over the time (in seconds). GFP-SofT n = 50 in 5 hyphae. mCherry-ChsB n = 50 in 6 hyphae. **(e)** Localization of GFP-SofT (depicted in green) and mCherry-ChsB (depicted in magenta) during hyphal tip growth. Numbers indicate the time in minutes. **(f)** Localization of ChsB-mCherry at the hyphal tip of the *A. flagrans* Δ*sofT*-mutant strain. The relative fluorescent intensity at the hyphal tip (y-axis, arbitrary units) was measured over the time course.

To understand how the signal oscillations in a single hypha (monologue) might transit into the anti-phasic, cell-to-cell dialogue once a partner cell appears in the vicinity, we constructed a mathematical model. Specifically, we extended an existing model of cell-to-cell communication based on anti-phasic oscillations to explicitly account for the uptake of signaling components from the surrounding media (8). The concentration of signaling components in the immediate proximity to the cell wall of cell 1 and cell 2 is represented by two compartments, *Z*_1_ and *Z*_2_, respectively **(Fig. 2a)**. As a signaling component is taken up into a cell, activating components (modeled as compartments *A*_1_ and *A*_2_) become more concentrated in this cell. These activating components, in turn, stimulate the docking of cytoplasmic signaling components (*X*_1_ and *X*_2_) to the inside of the cell membrane (*Y*_1_ and *Y*_2_). The release of membrane-docked vesicles to the extracellular space is initially blocked by high levels of activator (*A*_1_ and *A*_2_). Secretion ultimately occurs when some of the activating components are converted into an auto-inhibitory component (*I*_1_ and *I*_2_), reducing levels of activating components, thereby allowing secretion of signaling components into the extracellular compartments (*Z*_1_ and *Z*_2_). In simulations containing only a single cell, similar to the experimental results monitoring the components in a single hypha, oscillations in these different components could be observed **(Fig. S1)**. When placing two cells in the simulation at different distances (*d*) from each other, oscillations of these two cells appear uncoordinated at long distances (monologue), and anti-phasic at short distances (dialogue) **(Fig. 2b-d)**. The role of the signaling component in the dialogue at a short distance can be seen in our simulations: high extra-cellular concentrations (*Z*_1_ and *Z*_2_) are attained only briefly, when one cell has secreted the component, and the other cell has not yet taken it up again **(Fig. 2c)**. This back-and-forth exchange of signaling component via secretion into and uptake from the extracellular space represents a physically credible mechanism to establish the cell-to-cell dialogue at a short distance. When we implement a simulation that mimics the growth of two hyphae towards each other in the form of a distance that decreases over time, a transitory phase where the uncoordinated oscillations (two monologues) become mutually entrained into anti-phasic oscillations (dialogue) can be seen **(Fig. 2e)**. As seen in these simulation results, the same regulatory mechanism, based on a signaling component that is exchanged via the extracellular space, can explain the transition from single cell oscillatory growth to anti-phasic synchronization between two approaching cells. Crucially, these simulations also obviate a transitory phase, during which both cells’ dynamics slow down, just before entering into the dialogue and speeding up again **(Fig. 2e)**. This “critical-slowing down” is typical for dynamic systems when transitioning into a qualitatively different type of behavior (1, 23).

**Fig. 2:**
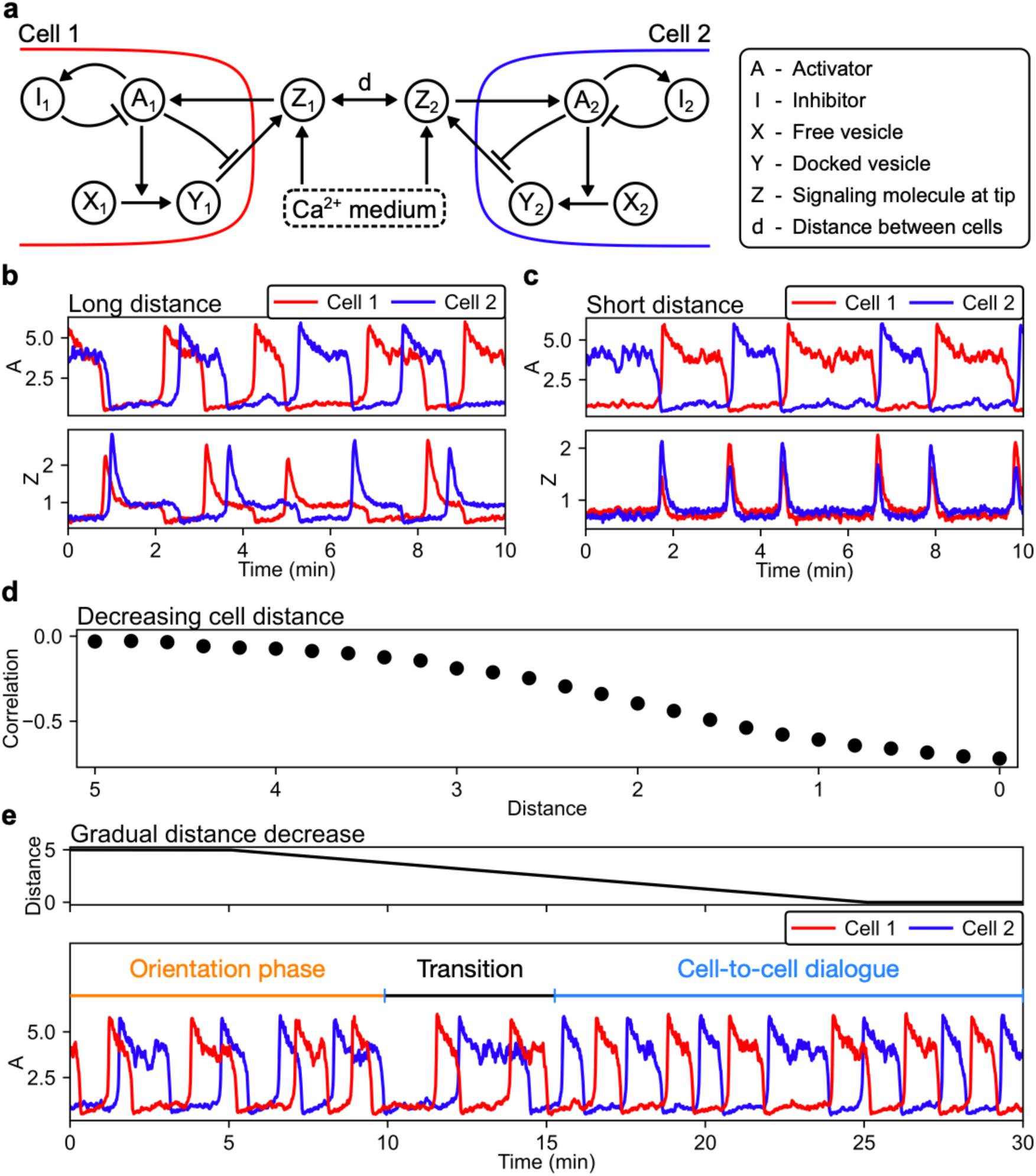
A mathematical model demonstrates the emergence of the temporal synchronization of two hyphae approaching each other. **(a)** In the model, each hyphal cell contains an excitatory system with an activator (*A*_1,2_) and inhibitor (*I*_1,2_). This excitatory system is triggered by extracellular signaling molecules (*Z*_1,2_) and, in turn, regulates the docking and subsequent release of vesicles (*X*_1,2_ and *Y*_1,2_), which contain the signaling protein, into the extracellular space. **(b)** At long distances (here, *d* = 10), the cells operate as independent oscillator units. **(c)** At short distances (here, *d* = 0), cells synchronize and oscillate in anti-phase. **(d)** Decreasing the distance between the two cells increases the magnitude of the anti-correlation (negative Pearson correlation coefficient) between the activator concentrations of the individual cells. **(e)** Upon continuous reduction of the distance between the cells, anti-synchrony emerges (noticeable at around 15 minutes).

To experimentally assess the transition from two “monologues” to a coordinated “dialogue”, we monitored SofT in two approaching hyphae. Initially, the oscillations in the two hyphae appeared uncoordinated (orientation phase) but then transitioned to anti-phasic oscillations characteristic of the cell-to-cell dialogue (**Fig. 3a, b, Movie S4**). The frequency of SofT oscillations during the cell-to-cell dialogue (mean period of 101 ± 30 (SD) seconds, n=50 in 5 hyphae) was comparable to the frequency in single hyphae **(Fig. 3c)**. As expected from our simulations of a dynamic transition into coordinated oscillations, oscillations in both hyphae slowed down during the transitory phase that precedes the coordinated cell-to-cell dialogue **(Fig. 3b)**. The underlying hypothesis that the same oscillation mechanism acts during the monologue- and dialogue-type dynamics is further substantiated by the observation that, similar to single hyphal growth, not only the SofT protein but also ChsB showed oscillating recruitment to the tips of interacting cells (mean period of 103 ± 26 (SD) seconds (n=105 in 11 hyphae)) **(Fig. S2a, Fig. 3c)**.

**Fig. 3:**
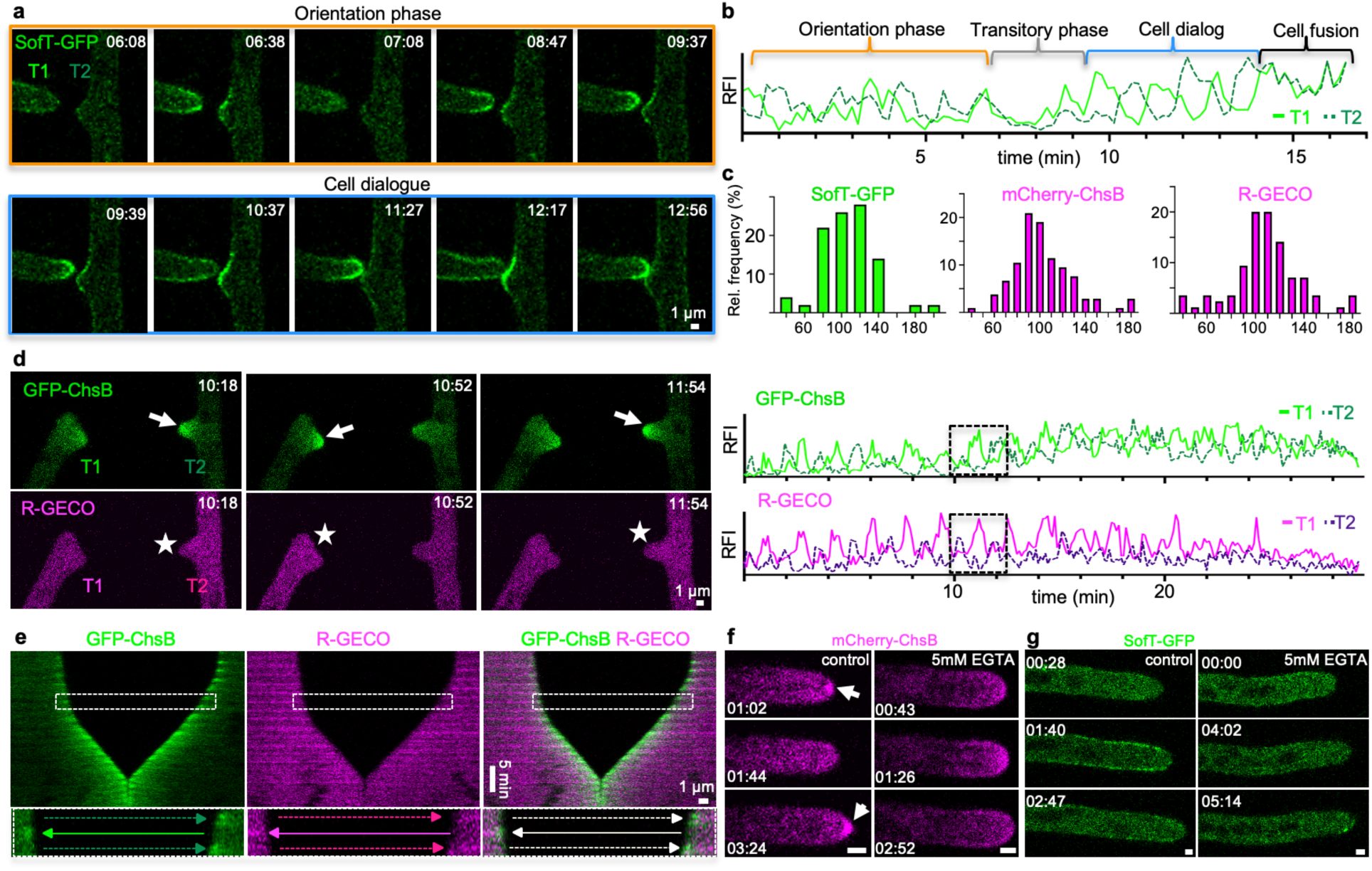
The stepwise extension of hyphae is coordinated during the cell dialogue stage of hyphal fusion. **(a, b)** Time course of SofT-GFP during a hyphal fusion event. Selective micrographs of the time course are shown and are divided into an orientation phase (orange frame) and cell dialogue phase (blue frame). An area of 3×3 pixels at each hyphal apex was used to measure the fluorescent intensity depicted in (b) (y-axis, arbitrary units). **(c)** The interval between two peaks of SofT-GFP, mCherry-ChsB, or R-GECO at each hyphal tip during the cell dialogue was counted and the distribution is depicted as relative frequency (y-axis) over the time (in seconds). SofT-GFP n = 50 in 3 hyphae. mCherry-ChsB n = 105 in 11 hyphae. R-GECO n = 85 in 12 hyphae. **(d)** Maximum-intensity projection of a time course of GFP-ChsB (depicted in green) and R-GECO (depicted in magenta) during a hyphal fusion event. Arrows indicate the localization of GFP-ChsB at hyphal tips. Stars indicate the fluorescent excitation of R-GECO in the presence of Ca^2+^ inside the hyphae. The RFI (y-axis) was measured of an area of 6×6 pixels at each hyphal apex over the time course (x-axis in minutes, y-axis, arbitrary units). The boxed area of the selected micrographs represents the enlarged area in **(e). (e)** A kymograph was created of the time course in (d) drawing a line (pixel width 5) along the growth axis of both hyphae. The boxed area is enlarged and displays one cycle of the cell dialog. Full arrows (light green, magenta) indicate the localization of the fusion proteins in the left hypha (T1). Dashed arrows (dark green, purple) indicate the localization of the fusion proteins in the right hypha (T2). **(f)** Compared to the control (LNA containing 1 μM CaCl_2_), mCherry-ChsB did not accumulate at the Spitzenkörper on LNA containing 5mM EGTA. **(g)** Compared to the control, GFP-SofT did not accumulate at hyphal tips on LNA containing 5 mM EGTA. Scale bars in (f, g) depict 1μm.

A central assumption underlying the cell-to-cell dialogue in our model is that hyphae communicate by passing back and forth a shared signaling component, *Z*. Considering the central role of Ca^2+^ in many types of cellular excitatory dynamics, and previous results that suggest that Ca^2+^ is involved in the cell-to-cell dialogue (24-27), we monitored intracellular Ca^2+^ concentrations using the genetically encoded fluorescent reporter R-GECO. The fluorescent signals showed robust oscillations that were coordinated between two approaching hyphae **(Fig. S2b, Movie S5)**(28). The mean oscillation period of 109 ± 26 (SD) seconds (n=85 in 12 hyphae) was comparable to the other markers during the cell-to-cell dialogue **(Fig. 3c)**. Indeed, simultaneous visualization of R-GECO and GFP-ChsB showed synchronized oscillation with similar periods, indicating the anti-phasic oscillations of growth during the chemotropic interaction of the two hyphae **(Fig. 3d, e, Movie S6)**. This phenomenon was coordinated in anti-phase in interacting hyphae. These results indicate that signaling and growth during the cell-to-cell dialogue are highly synchronized and probably mediated by the uptake of Ca^2+^.

To directly test the role of Ca^2+^, we depleted Ca^2+^ from the media by adding the Ca^2+^ chelating agent EGTA to LNA containing 1 μM CaCl_2_. At an EGTA concentration of 5 mM, cell-fusion events were never observed after incubation for 16 up to 72 hours **(Fig. S3a)**. In addition, hyphal growth of *A. flagrans* was reduced, but germination of spores was unaffected. Fluorescence of R-GECO was not detectable in these Ca^2+^-depleted conditions, indicating that intracellular Ca^2+^ oscillations indeed rely on extracellular Ca^2+^ **(Fig. S3b)**. mCherry-ChsB still localized at hyphal tips, however, no obvious oscillating dynamic recruitment was observed **(Fig. 3f)**. mCherry-ChsB localized directly to the plasma membrane at the tip, without prior accumulation at the Spitzenkörper, indicating the importance of Ca^2+^ for well-regulated pulsatile secretion. The localization of GFP-SofT to hyphal tips was abolished after the addition of 5 mM EGTA, indicating Ca^2+^-dependent recruitment to the plasma membrane **(Fig. 3g)**. In line with these experimental results, reducing the external media concentration of signaling components in our simulations also abolished oscillations **(Fig. S4)**. Taken together, these results show that extracellular Ca^2+^ is essential in the signaling mechanism that underlies the monologue implicated in pulsatile cell extension as well as the synchronization of oscillations into a cell-to-cell dialogue **(Fig. 4)**. In the future, it will be interesting to study how other fusion-related proteins might be involved in the translation of the increase in intracellular Ca^2+^-concentrations and how this change relates to the secretion of a chemo-attractive signal molecule. The growing tip of filamentous fungi is an example of apical-growing cells and therefore our results may have an impact on the understanding of other highly polarized cells such as plant pollen tubes or root hairs, or axons and dendrites (1).

**Fig. 4:**
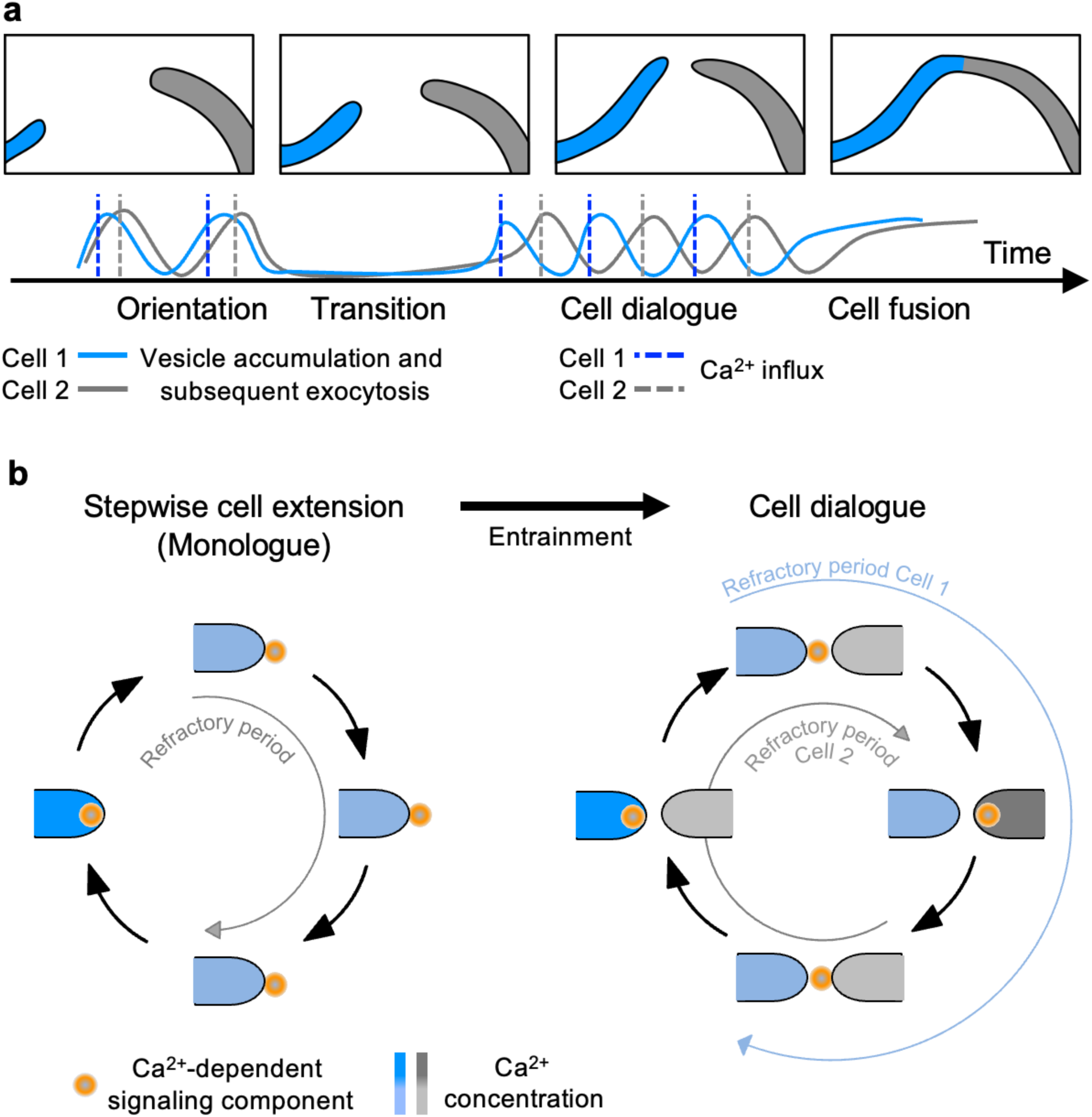
Scheme of the cell-to-cell dialogue. **(a)** Cell fusion of two hyphal cells is mediated by the synchronization of oscillatory growth. Vesicles needed for hyphal growth and communication accumulate at the hyphal tip and are released to the surroundings upon an influx of Ca^2+.^ If two cells are in close proximity, the uncoordinated, stepwise growth shifts after transitory entrainment to a synchronized anti-phasic cell dialogue and subsequent cell fusion. **(b)** During stepwise cell extension, a Ca^2+^-dependent signaling component is released with a refractory period to prevent self-stimulation. Entrainment of two cells in close proximity initiates a cell dialogue mediated by a so far unknown Ca^2+^-dependent signaling component. Refractory periods after secretion prevent self-stimulation.

## Methods

### Strains and culture conditions

*A. flagrans* strains (all derived from CBS349.94) were cultivated at 28 °C on potato dextrose agar (PDA, 2.4% potato dextrose broth and 2% agar, Carl Roth). All fungal strains used in the study are listed in **Table 1**. Protoplast transformation was performed as described in (29). Chemically competent *Escherichia coli* Top10 cells were used for plasmid cloning.

**Table 1:** *A. flagrans* strains used in this study.

| Strain number | Genotype | Origin |
| --- | --- | --- |
| CBS 349.94 | Wild type | (29) |
| sVW16 | *tubA(p)::lifeact::GFP::gluC(t); trpC(p)::hph::trpC(t)* | (31) |
| sVW01 | *sofT::hph (ΔsofT)* | (29) |
| sVW29 | *sofT(p)::sofT::GFP::sofT(t); trpC(p)::hph::trpC(t)* | This study |
| sVW51 | *tubA(p)::mCherry::chsB::chsB(t); gpdA(p)::neo::trpC(t)* | This study |
| sVW52 | *sofT(p)::GFP::sofT;*<br>*tubA(p)::mCherry::chsB::chsB(t);trpC(p)::hph::trpC(t);*<br>*gpdA(p)::neo::trpC(t)* | This study |
| sVW65 | *h2b(p)::r-GECO::gluC(t); gpdA(p)::neo::trpC(t)* | This study |
| sVW69 | *h2b(p)::r-GECO::gluC(t);tubA(p)::GFP::chsB::chsB(t);*<br>*gpdA(p)::neo::trpC(t); trpC(p)::hph::trpC(t)* | This study |
| sVW75 | *sofT::hph (ΔsofT); tubA(p)::mCherry::chsB::chsB(t);*<br>*gpdA(p)::neo::trpC(t); trpC(p)::hph::trpC(t)* | This study |

### Plasmid and strain construction

All plasmids and primers used in the study are listed in **Table 2 and 3**. Chemically competent *Escherichia coli* Top10 cells were used for plasmid cloning. *A. flagrans* SofT was tagged with GFP at the C-terminus and expressed under the native promoter at the *sofT* locus. A 1 kb region from the 3’-end of the sofT *open reading frame* (orf) and 0.6 kb of the terminator region were amplified by PCR from *A. flagrans* genomic DNA. A *GFP*::*hph* cassette was amplified from pNH57 and all three fragments were subsequently inserted into the linearized plasmid backbone pJET1.2 using Gibson assembly, resulting in plasmid pVW106. *A. flagrans* ChsB was tagged with either mCherry or GFP at the N-terminus and expressed under the alpha-tubulin *tubA*-promoter. The *chsB* orf and the 0.9 kb 3’ region were amplified by PCR from *A. flagrans* gDNA and inserted into the plasmid backbone pVW92 using Gibson assembly, resulting in the *tubA*-*mCherry*-*chsB* plasmid pVW118. For a GFP-tagged ChsB variant, the *mCherry* cassette of pVW118 was replaced by *GFP* using Gibson assembly, resulting in pVW132.

**Table 2:** Plasmids used in this study.

| Name | Description/Genotype | Reference |
| --- | --- | --- |
| pVW57 | Plasmid backbone containing *trpC(p)::hph::trpC(t)* | (31) |
| pVW92 | Plasmid backbone containing *gpdA(p)::neo::trpC(t)*<br>and *tubA(p)* | (13) |
| pNH57 | Plasmid containing *GFP::hph* | Nicole Wernet<br>(Karlsruhe) |
| pVW106 | *sofT(p)::sofT::GFP::sofT(t); trpC(p)::hph::trpC(t)* | This study |
| pVW118 | *tubA(p)::mCherry::chsB::chsB(t);gpdA(p)::neo::trpC(t)* | This study |
| pVW132 | *tubA(p)::GFP::chsB::chsB(t);*<br>*gpdA(p)::neo::trpC(t)* | This study |
| pVW120 | *tubA(p)::r-GECO::gluC(t); gpdA(p)::neo::trpC(t)* | This study |
| pVW125 | *h2b(p)::r-GECO::gluC(t); trpC(p)::hph::trpC(t)* | This study |
| pVW131 | *h2b(p)::r-GECO::trpC(t); trpC(p)::hph::trpC(t)* | This study |

**Table 3:**
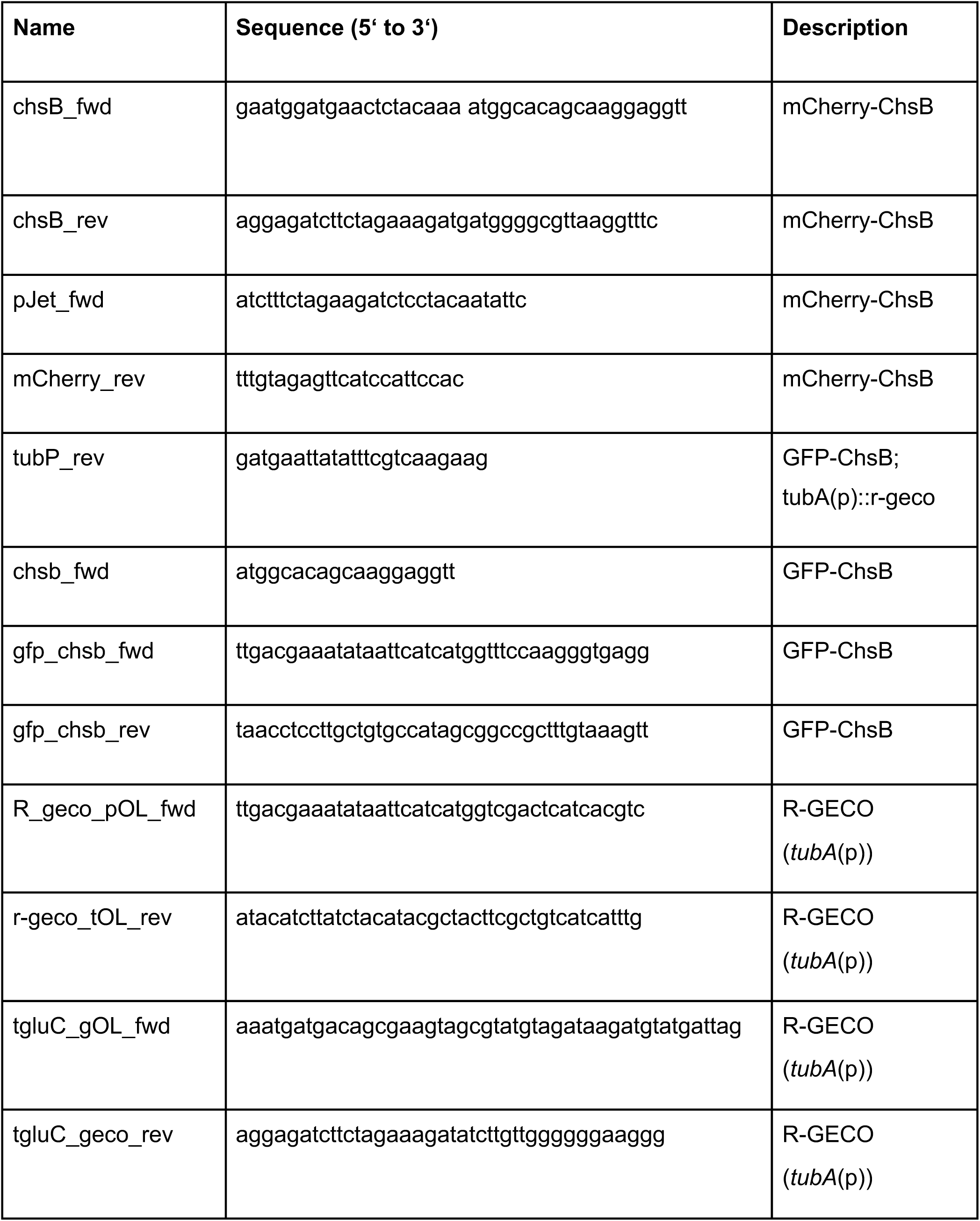
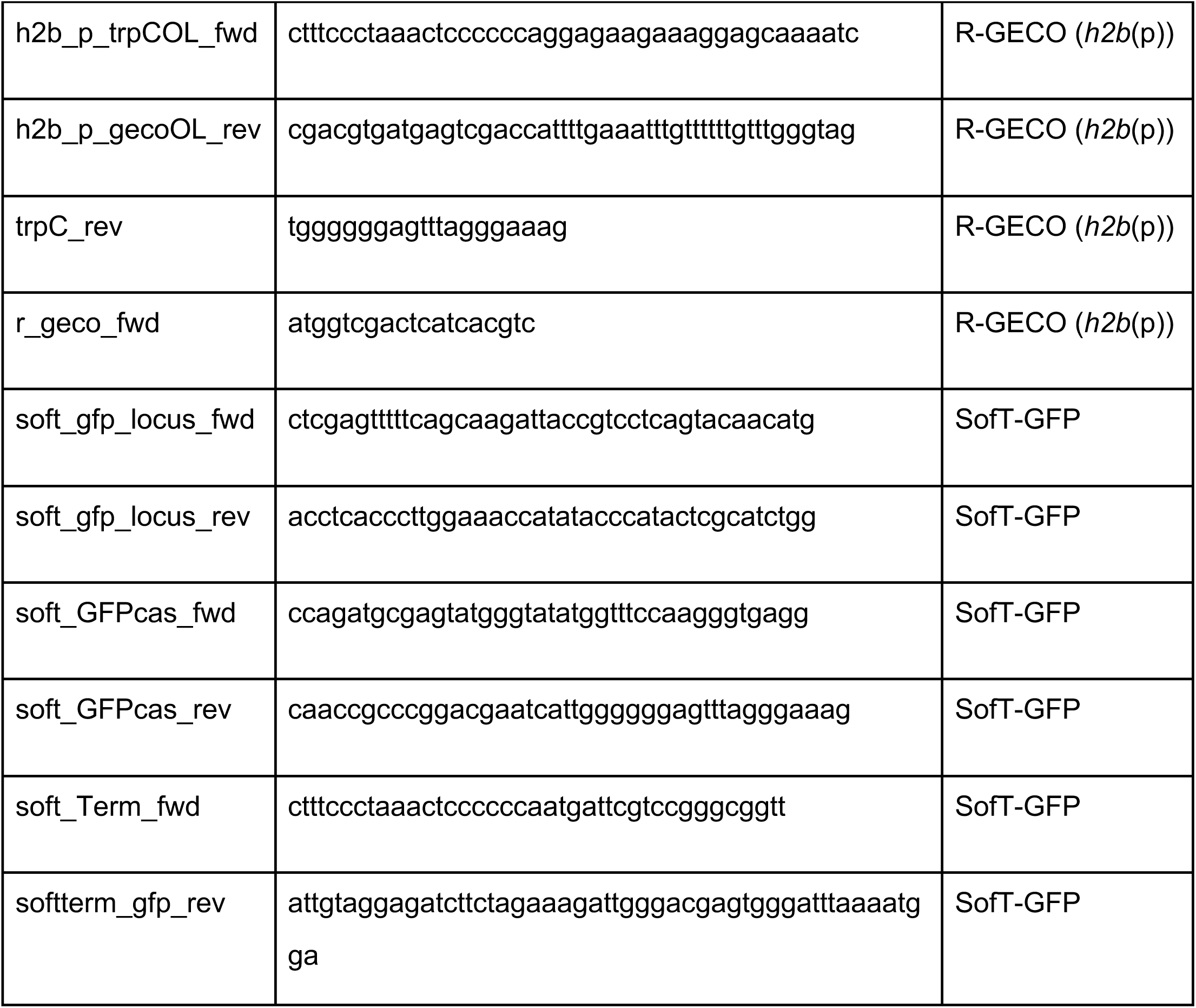
Oligonucleotides used in this study.

R-GECO was expressed under the histone *h2b*-promoter. The R-GECO sequence was amplified from gDNA of SNT162 (4) and cloned into plasmid pVW92, resulting in pVW120. Subsequently, the *tubA*-promoter was exchanged by the 1 kb sequence of the *h2b* promoter using Gibson assembly, resulting in plasmid pVW125. For co-localization experiments, the *h2b*(p)-*R-GECO*-fragment was cloned into pVW57, resulting in pVW131.

### Microscopy

For microscopy, fungal strains were inoculated on thin low-nutrient agar (1g/l KCl, 0.2 g MgSO4 - 7H_2_O, 0.4 mg MnSO_4_ - 4H_2_O, 0.88 mg ZnSO_4_ - 7H_2_O, 3 mg FeCl_3_ - 6H_2_O, 15 g agar, pH 5.5) slides supplemented with 1μM CaCl_2_. For calcium chelating experiments EGTA (stock solution 0.5 M) was added at a final concentration of 5 mM. Around 1×10^4^ *A. flagrans* spores were incubated on a 2×2cm agar pad at 28 °C in darkness for 12 – 72 h.

Live cell imaging was performed using a confocal microscope (LSM900, Carl Zeiss) with a 63x NA 1.4 oil objective lens (DIC M27). Time series were acquired with a gallium arsenide phosphide photomultiplier tube (GaAsP-PMT) detector, 488 or 561 nm excitation lasers were used. Image processing and analysis were performed in FIJI (30) and ZEN Blue. Datasets were analyzed in GraphPad Prism. Kymographs were generated with the FIJI Multi Kymograph tool using line widths 5. The dynamics of the fluorescent intensity over time were measured in each frame with a circular selection at the respective hyphal tip, specified in each figure legend. The interval between two accumulating peaks at hyphal tips was counted in ZEN Blue.

All raw images and data files resulting from the analyses are available at Zenodo (https://zenodo.org/record/6830734#.Ys_KmS2w1TY).

### Mathematical model

Our model is based on the previous model of the dialogue-like communication between two Neurospora cells growing towards each other (8). The original model consists of eight ordinary differential equations, four representing each cell. Those equations represent an excitatory system that coordinates the docking and release of vesicles with chemoattractants. We extended this model by explicitly modeling the dynamics of the chemoattractants in the extracellular space at the tips of the individual cells. This is modeled by two coupled differential equations that represent the concentration of the signaling molecules in the proximity of the corresponding cells. Those two equations are linked with a coupling whose magnitude represents the coupling strength. The model was implemented in the Julia Programming Language as a set of stochastic differential equations and simulated using the Euler-Maruyama method with integration step dt=0.001. The complete set of equations and parameter values can be found in the Supplementary Information. The simulation script is also available at [https://github.com/vkumpost/cell-dialog].

## Acknowledgement

We are thankful to the group of Prof. Dr. André Fleißner (University of Braunschweig) for fruitful discussions and Prof. Dr. Sylvia Erhardt (KIT) for the opportunity and the help to use the Zeiss LSM900 microscope. Valentin Wernet was funded by the German Federal Environmental Foundation (DBU). VK was funded by the Helmholtz Information and Data Science School for Health (HIDSS4Health). RM and LH were funded by the Helmholtz Program Natural, Artificial, and Cognitive Information Processing (NACIP).

## Author contributions

**Valentin Wernet:** Investigation, validation, writing - original draft, writing – review & editing, visualization. **Vojtech Kumpost:** Investigation, visualization. **Ralf Mikut:** writing – review & editing, supervision, funding acquisition. **Lennart Hilbert:** Investigation, supervision, writing – review & editing. **Reinhard Fischer:** Supervision, writing – review & editing, project administration, funding acquisition.

## Additional information

Supplementary information is available online. Correspondence and requests for material should be addressed to R.F.

## Competing interests

The authors declare no competing interests.

## Figures

**Fig. S1:**
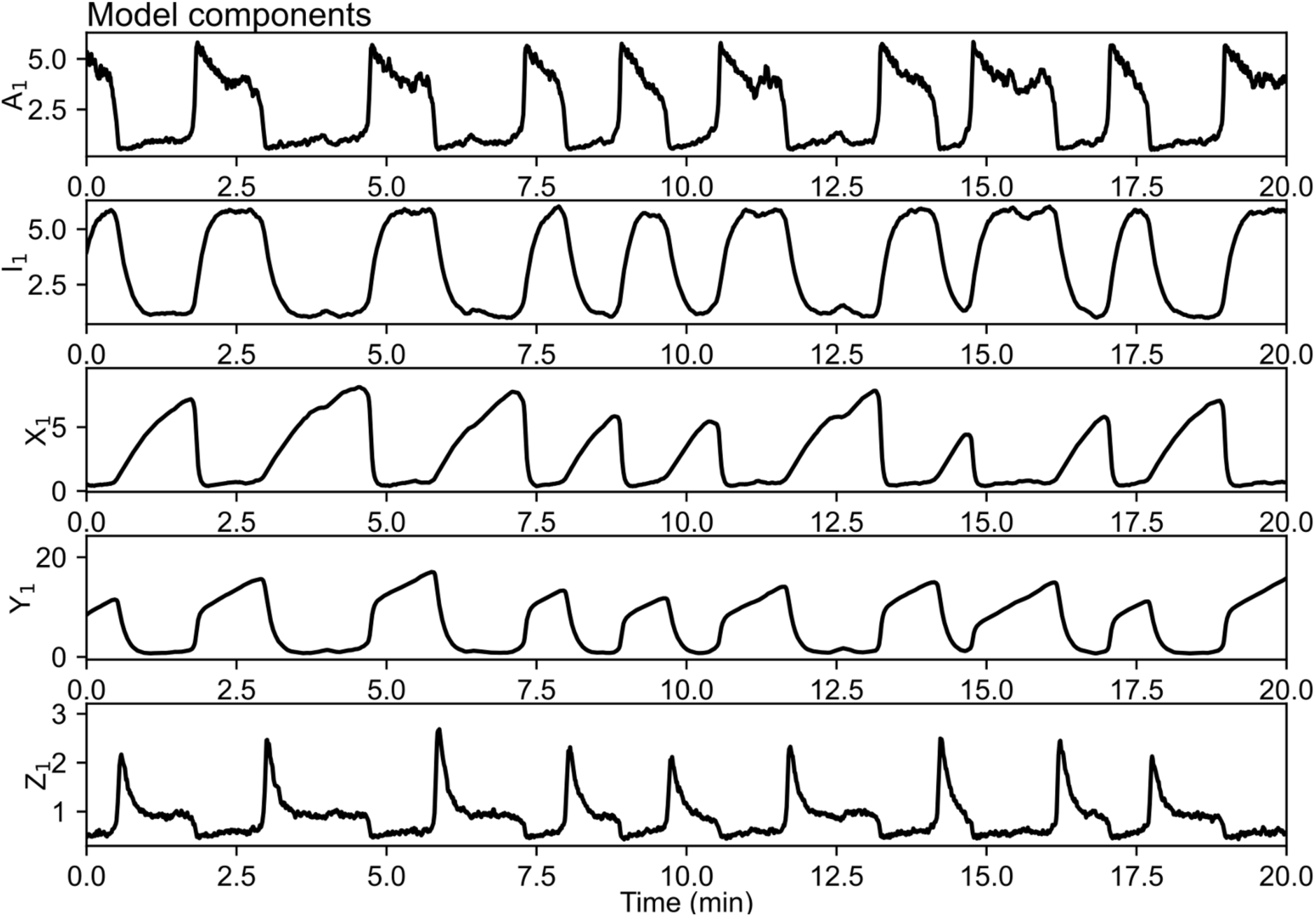
Example simulation containing a single cell to illustrate the oscillatory dynamics of all model variables. To illustrate the time course of all model variables, a simulation with a single cell was implemented by setting the cell distance to infinity (*d* = ∞).

**Fig. S2:**
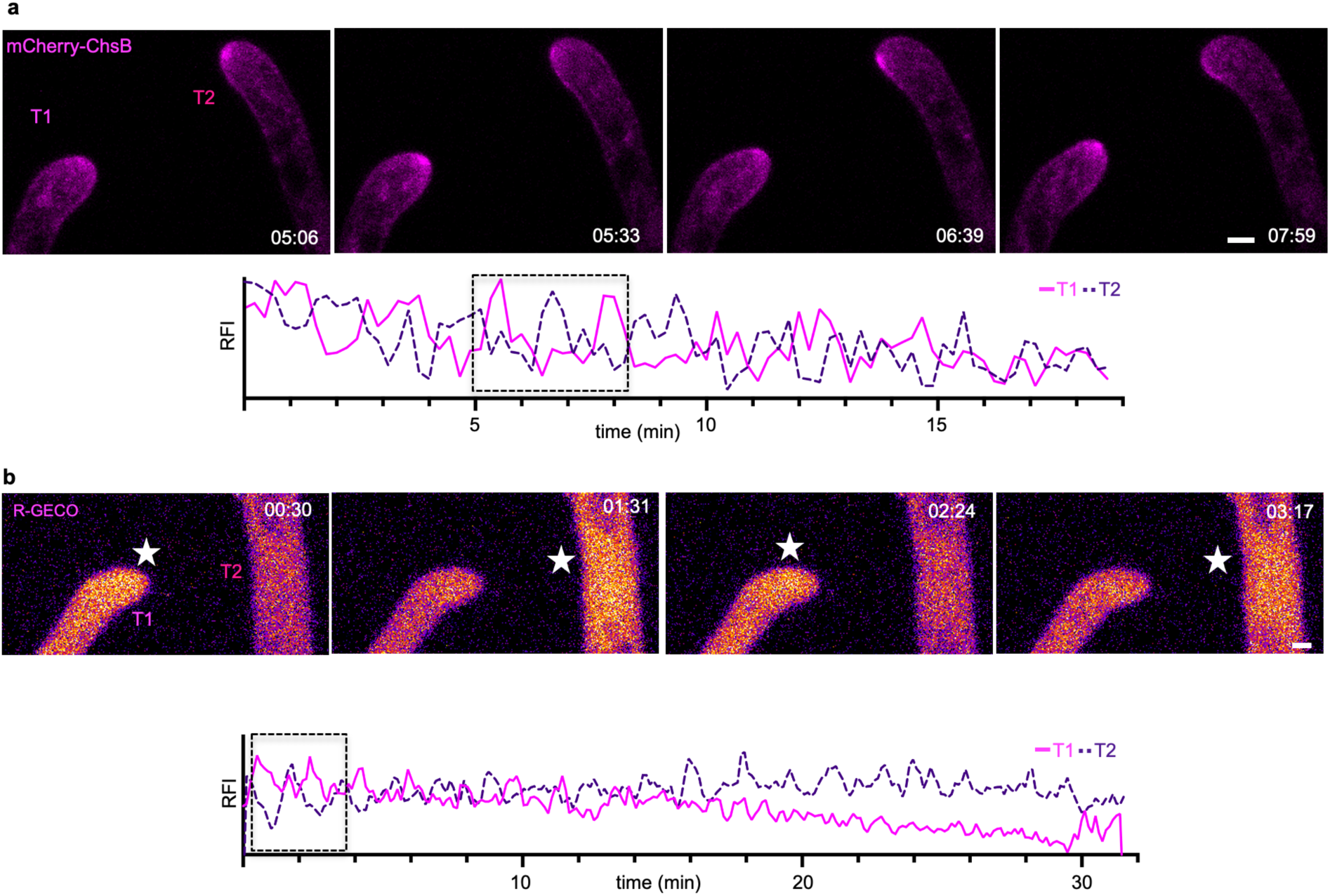
The oscillating extension of hyphae is synchronized during hyphal fusion in *A. flagrans*. **(a)** Time course of mCherry-ChsB during a hyphal fusion event. A maximum-intensity projection was generated from the time-lapse sequence. It was further bleach-corrected using the bleach correction plugin (Correction Method: Simple Ratio) of Fiji. The RFI of an area (y-axis, arbitrary units) of 6×6 pixel was measured at each hyphal apex over time (x-axis in minutes) to generate the graph. The boxed area depicts the selected micrographs of the time course. **(b)** Maximum-intensity projection of a time course of R-GECO during a hyphal fusion event. Stars indicate the fluorescent excitation of R-GECO in the presence of Ca^2+^ inside the hyphae. The changes in fluorescent intensity are color-coded as Fire LUT using Fiji, depicting high pixel values as white and yellow color, and low pixel values as blue and magenta. The RFI (y-axis, arbitrary units) was measured at each hyphal tip over the time course (x-axis in minutes). The RFI of an area of 12×12 pixels was measured at each hyphal tip to generate the graph. Scale bars in (a, b) depict 1μm. The boxed area depicts the selected micrographs of the time course.

**Fig. S3:**
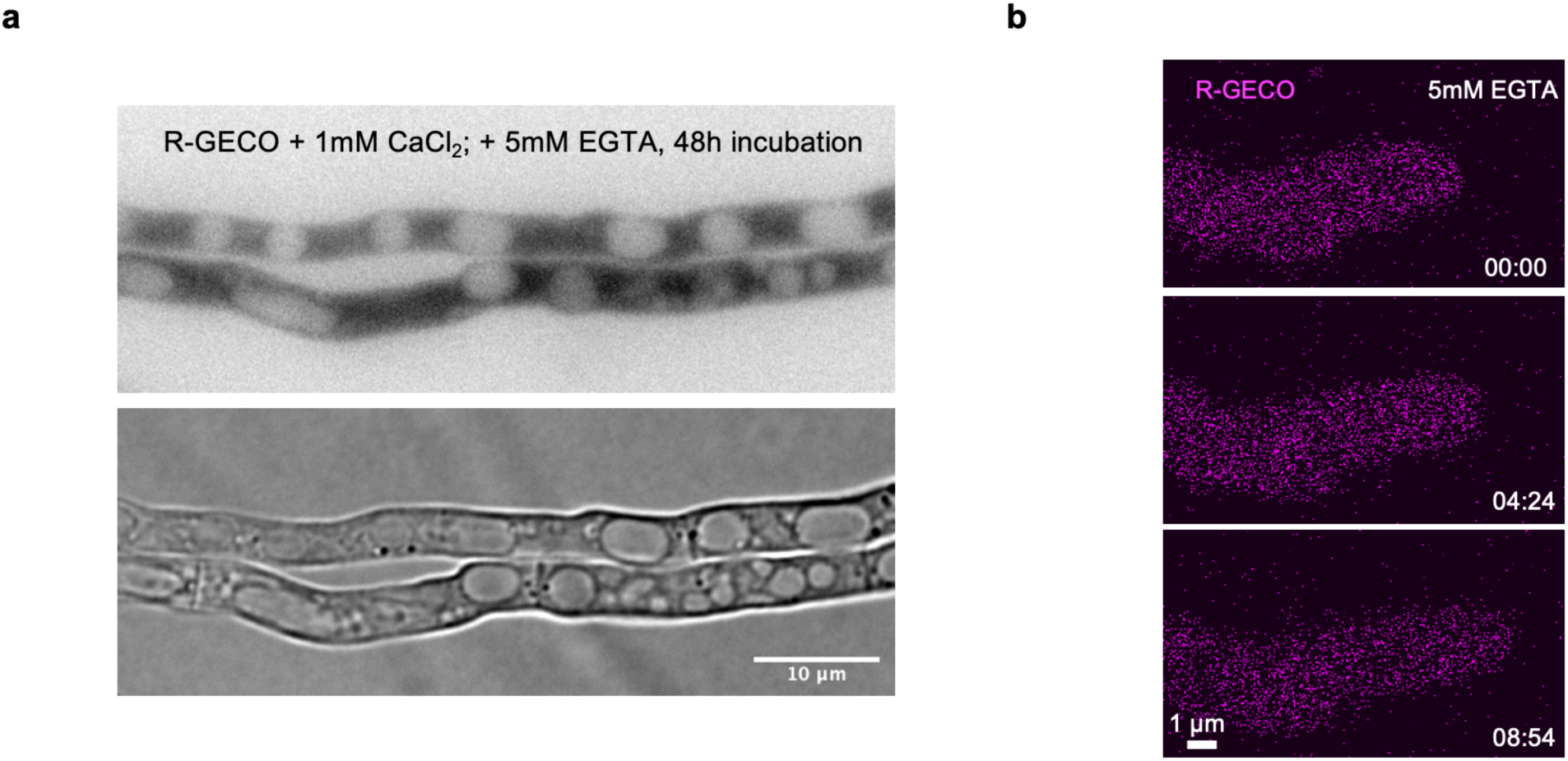
Extracellular Ca^2+^ is necessary for hyphal fusion and pulse-like exocytosis. **(a)** Hyphae showed close physical contact without hyphal fusion after incubation with the Ca^2+^ chelating agent EGTA (5mM). **(b)** No pulses of R-GECO were observed in hyphae on LNA containing 5mM EGTA.

**Fig. S4:**
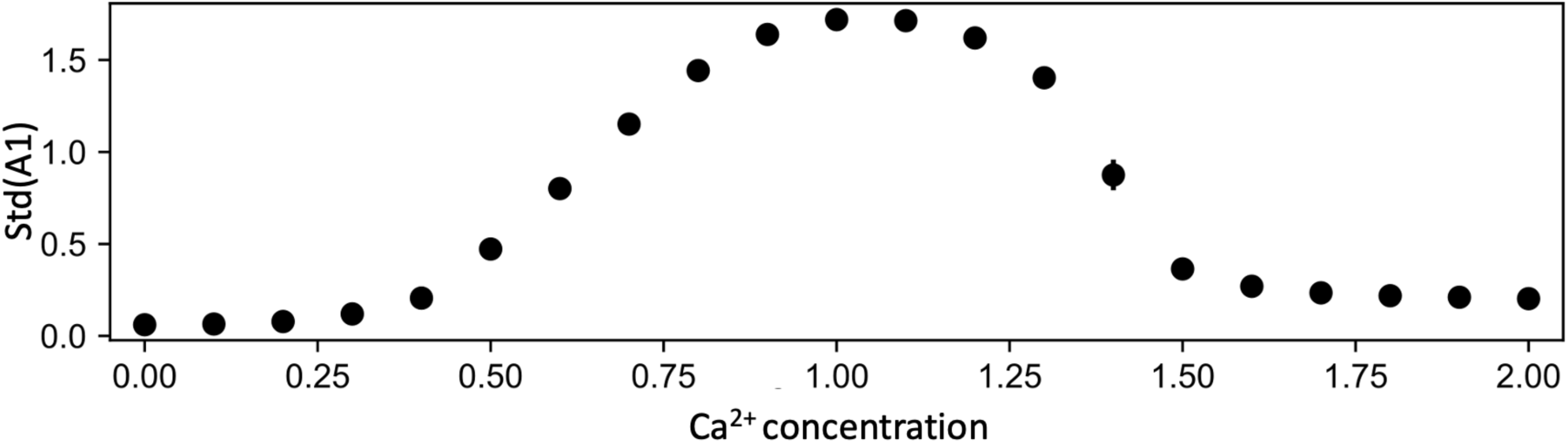
Sufficient concentration of Ca^2+^ is necessary to produce oscillations. In a simulation containing only a single cell, at a Ca^2+^ concentration of 0 (*C*_*media*_= 0), no oscillations are observed. Around a Ca^2+^ concentration of 1 (*C*_*media*_= 1), the cell generates regular oscillations (as shown in Fig. S1), evidenced by an increasing standard deviation of the activator level (Std(*A*_1_)). Increasing the Ca^2+^ concentration further leads again to the gradual loss of oscillatory behavior. The standard deviation was calculated over a simulation that represented 800 minutes of cellular dynamics.

**Movie S1:** Localization of GFP-SofT during hyphal tip growth.

**Movie S2**: Localization of mCherry-ChsB during hyphal tip growth.

**Movie S3**: Co-localization of GFP-SofT (depicted in green) and mCherry-ChsB (depicted in magenta) during hyphal tip growth.

**Movie S4:** Time course of GFP-SofT during a hyphal fusion event.

**Movie S5:** Time course of R-GECO during a hyphal fusion event.

**Movie S6:** Time course of GFP-ChsB (depicted in green) and R-GECO (depicted in magenta) during a hyphal fusion event.

## Model Supplementary Information

The model was implemented in the form of stochastic differential equations as

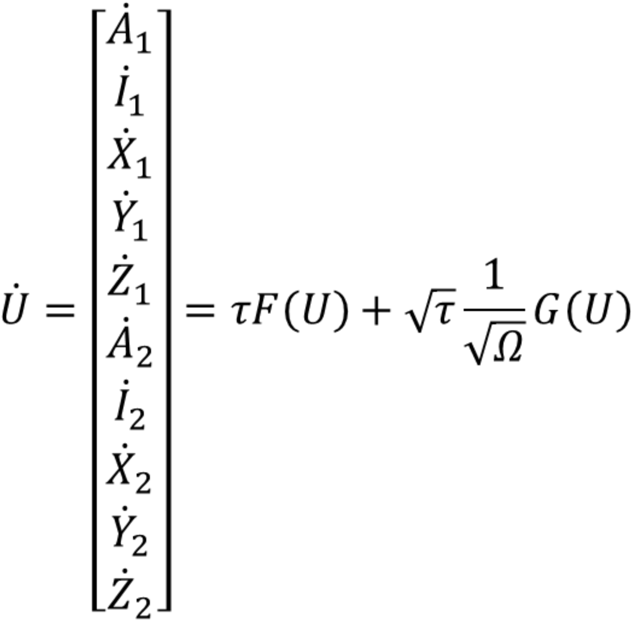

where U is a vector of state variables that represent activator (A), inhibitor (I), free vesicle (X), docked vesicle (Y) and concentration of the extracellular signaling molecule at the tip of the cell (Z). The index (1, 2) indicates the specific cell. τ is a time-scaling constant to adjust the time scale without the loss of dynamics. Ω is the system size and controls the level of noise in the system. The drift function (F) reads

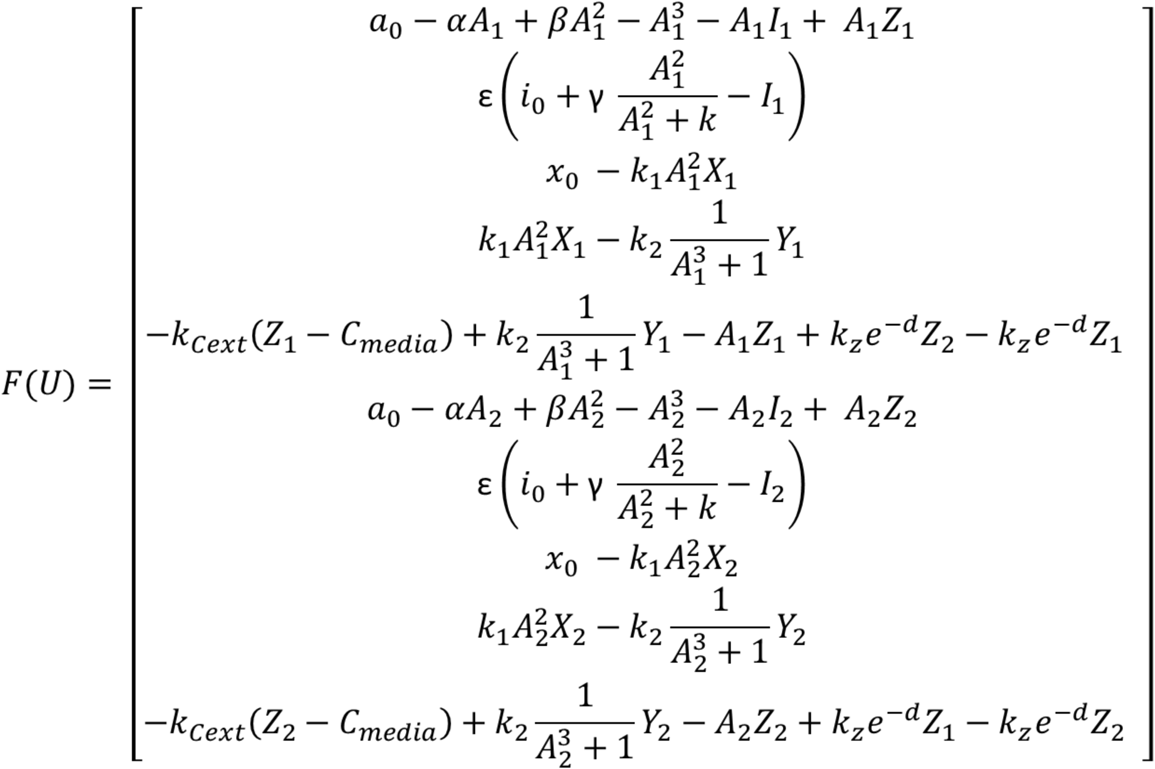

The noise function (G) is

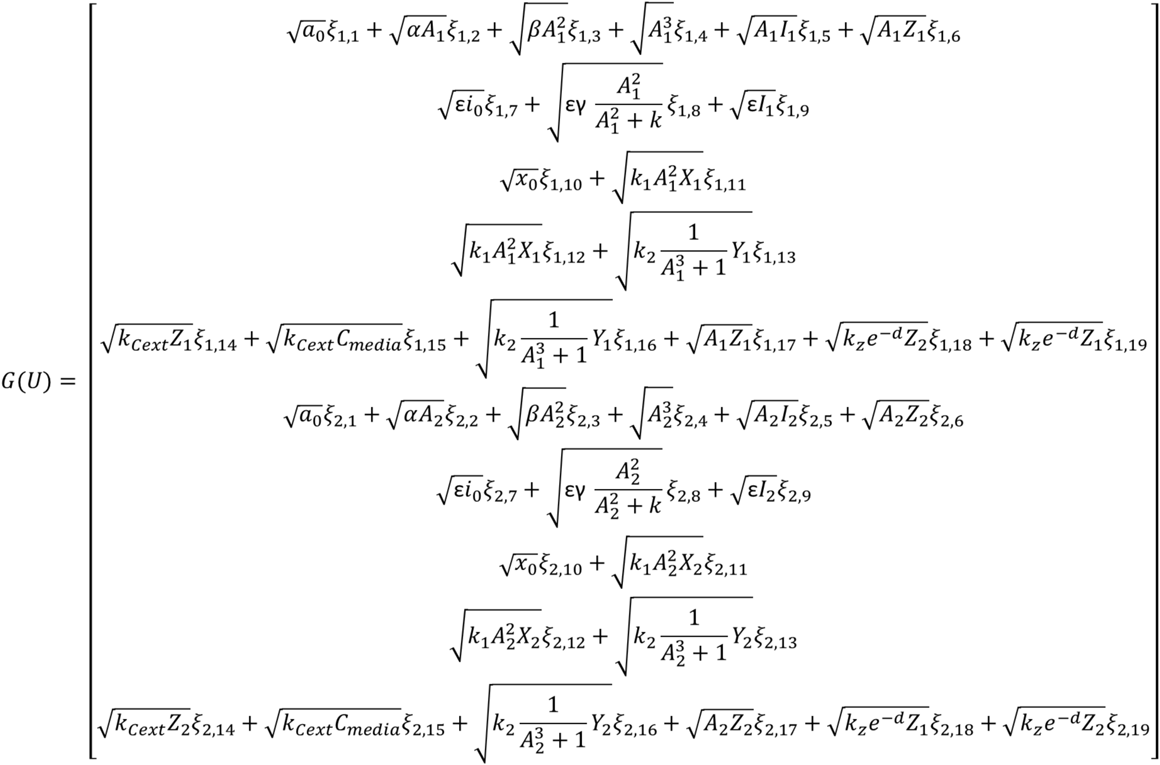

where ξ are independent Wiener processes. The results presented in the manuscript were obtained for parameter values τ = 8.48, ε = 0.55, α = 12.4, β = 8.05, γ = 8, k = 6, a_0_ = 5.6, i_0_ = 0.1, x_0_ = 1.0, k1 = 0.1, k2 = 1.0, kz = 10.0, kcext = 5, Cmedia = 1, d = 10 (long distance), d = 0 (short distance), Ω = 1000. The initial conditions for the simulations were set to U(t=0) = [0.7065, 0.7145, 0, 0, 1.0, 0.7065, 0.7145, 0, 0, 1.0]. To minimize the effect of the initial conditions the model was run for 200 minutes before any other analysis was performed.

## Notes

### Competing Interest Statement

The authors have declared no competing interest.

